# Stable, intronic RNAs explain preservation of introns in *Cyanidioschyzon merolae*

**DOI:** 10.1101/2025.12.01.691491

**Authors:** Patrick Geertz, Viktor Slat, Alison Ziesel, Maryam Ghaffarzadeh, Breanne Hatfield, Hosna Jabbari, Ben Montpetit, Martha Stark, Stephen Rader

## Abstract

Despite recent work identifying functional roles for introns, we lack a broad understanding of why some introns are preserved while others are removed. Here, we use the thermophilic red alga, *Cyanidioschyzon merolae*, as a model to investigate why only 39 of the approximately 2000 ancestral introns were preserved in this lineage. We observe that 23 of the 39 introns encode stable RNAs, 11 of which represent a novel class of non-coding RNA (ncRNA), which we call stable intronically-encoded RNAs (sieRNAs). These novel ncRNAs are expressed constitutively under normal growth conditions and are conserved in other extremophilic algae. One sieRNA, *Q270*, is predicted to stabilize a chloroplast polycistronic transcript encoding 17 ribosomal protein genes through direct antisense base pairing. Strikingly, all sieRNAs are polyadenylated under heat stress with long tails ranging from 50-200 nucleotides long, linking them to a potential heat stress response that may be critical for heat adaptation. Furthermore, DMS-MaP chemical probing revealed that some sieRNAs contain three-way junctions, a common RNA regulatory element, while others undergo accessibility shifts between *in vivo* and *in vitro* conditions, indicative of cellular interactions. Our findings suggest that introns in *C. merolae* are preserved to encode ncRNAs and suggest that introns may serve as hosts for regulatory RNAs across eukaryotes.

## BACKGROUND

Once considered “junk” sequences, introns are now recognized as important RNA features for cell survival (1,2). During pre-mRNA processing, they play an important role in regulating alternative splicing, while excised full-length introns can help cells survive under certain stress conditions (3,4). Furthermore, introns can be processed into various types of non-coding RNAs (ncRNAs) including small nucleolar RNAs (snoRNAs), long non-coding RNAs (lncRNAs), microRNAs, and stable intronic sequence RNAs (sisRNAs) (5–7). The functions of many ncRNAs have been previously annotated and are often involved in regulating cell development and cellular adaptation to stress conditions (8,9). However, it is still poorly understood why some introns are removed from the genome over evolutionary time, while others persist. This suggests that an effective system is required to assess the preservation of introns and to strengthen our understanding of their potential roles.

*Cyanidioschyzon merolae* is a unicellular red alga found in volcanic hot springs (10,11). Commonly studied for its simple cellular architecture, *C. merolae* also has a relatively low number of introns, making it an excellent model for studying whether they are functional. Recent work has identified that *C. merolae* contains 39 introns (Slat et al., manuscript in preparation), while other related red algal species are still relatively intron rich (12). This contrasting intron abundance suggests that *C. merolae* has lost a dramatic number of introns and raises the question of whether remaining introns are preserved due to their function, or if these introns will continue to be removed through evolutionary time.

In this work, we sought to characterize the remaining *C. merolae* introns genetically, biochemically, and computationally to explore their preservation. We identify a novel class of highly stable ncRNAs found in over half of the introns that are predicted to play various cellular roles under normal growth conditions and under stress. We suggest that these ncRNAs explain the preservation of introns in *C. merolae* through evolutionary time and reveal that introns may serve as a host for regulatory RNAs in eukaryotes.

## RESULTS

### Stable intronic RNAs identified in over half of *C. merolae* introns

As *C. merolae* has lost nearly 2000 introns through evolutionary time, the retention of only 39 introns suggests that they are important for *C. merolae* fitness. To explore whether the remaining introns in *C. merolae* are functional, we used long-read sequencing to search for excised, stable RNAs. In 23 of the 39 annotated introns, we identified intronic reads under normal growth conditions (Fig. 1A). These features are smaller than the full-length intron, lying between the 5’ splice site and the branch point. They range in size from 82 to 1298 nucleotides, and are observed in total RNA (ribodepleted) but not polyA+ RNA. To validate the RNA-seq data, we probed RNA blots for transcripts corresponding to the intronic features. Blots confirmed the presence of RNA species whose lengths matched the stable RNAs detected in RNA-seq data (Fig. 1B). Importantly, we were unable to detect intronic RNAs when probing for introns that did not have any features in the RNA-seq data (Fig. 1C, D), suggesting that this is not a universal phenomenon in *C. merolae* introns. It is possible that these RNAs represented splicing intermediates, as stable intronic RNAs can exist as lariats once excised through splicing (13). We noted, however, that they migrated on Northern blots consistently with their predicted sizes based on long-read sequencing data, suggesting linearity (Fig. 1B).

**Fig. 1.**
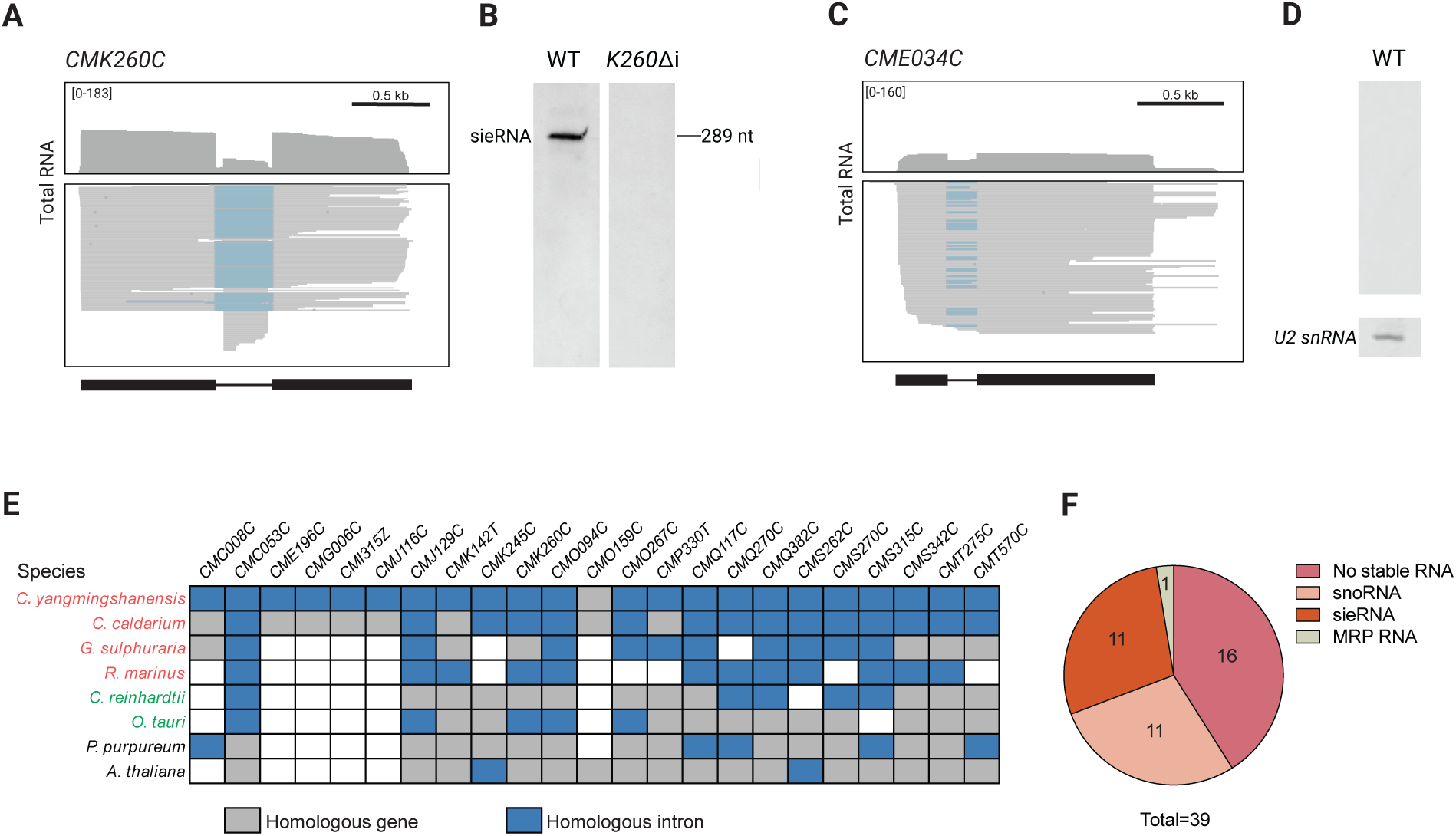
Stable RNAs in over half of *C. merolae* introns are evolutionary conserved. **A.** Detection of stable intronic RNA fragment in the *CMK260C* intron. PacBio ribo-depleted RNA-seq data with individual reads (lower box) and read trace (top box). Blue lines are absent introns in reads from spliced transcripts. Gene model (below): thick boxes, coding sequence; thin line, intron sequence. **B.** RNA blot probed for *CMK260C* intronic RNA using total RNA from wildtype (WT) and *CMK260C* intron deletion strain (*K260*Δi) cells. **C**. Absence of intronic feature in *CME034C* intron-containing gene as described in **A. D.** RNA blot probed for *CME034C* intron from WT cells. *n = 3* biological replicates were checked. U2 snRNA was probed as a loading control. **E.** Conservation of ncRNA-containing introns across select algae and plants. Thermophilic species are colored red, mesophilic green algae are colored green and plants are colored black. **F.** Identity of intron features in *C. merolae*.

Given that they represented a potentially novel class of ncRNA, we sought to determine whether intronic RNAs were conserved. We measured the conservation of introns harbouring ncRNAs across thermophilic red algae, green algae, and plants. ncRNA-containing introns were highly conserved in other thermophilic algae (11 to 22 of 23 introns) while green algae and plant species had fewer conserved introns (2 to 5 of 23 introns) (Fig. 1E). The higher degree of conservation among thermophilic algae suggests that these ncRNAs may provide an advantage for heat adaptation.

As some classes of ncRNA, such as small nucleolar RNAs (snoRNAs), have been reported in introns, we assessed whether these intronic RNAs belonged to previously annotated ncRNA classes using bioinformatic analyses. Of the 23 features, 11 corresponded to box C/D snoRNAs, and one was annotated as the RNase MRP RNA (Fig. 1F). These RNAs will be discussed separately in a future publication. This leaves 11 intronic RNAs that have not been previously characterized, which we call stable, intronically-encoded RNAs (sieRNAs).

### sieRNA deletion and growth phenotypes

As sieRNAs are expressed constitutively under normal conditions, we reasoned that they may serve an important function in cell growth and reproduction. To test this, we generated a series of intron-deletion mutants: strains lacking individual introns that contain sieRNAs, and controls without sieRNAs, and strains in which six (Δ6i) or ten introns (Δ10i) were deleted. We next compared their doubling times relative to wildtype (WT) using a plate-based growth assay (Fig. 2A). We expected that if sieRNAs were important for cell fitness, strains lacking sieRNA-containing introns would grow more slowly, and that removal of multiple sieRNAs would produce more dramatic growth defects. Notably, we only measured slower growth in *Q270*Δi strain, Δ6i and Δ10i relative to WT. While the growth defect in these strains could result from sieRNA loss, intronic sequences can also affect splicing efficiency of their host genes, leading to variable gene expression (13,14). To distinguish between these possibilities, we tested whether the expression of host genes changed when the intron was deleted using RNA-seq. Given that all introns in the Δ6i strain, including the *Q270* intron, are deleted in the Δ10i strain, we reasoned that expression changes in Δ10i would be representative of those observed in the Δ6i and *Q270*Δi strains. We observed that all intron-deleted host genes except *Q270C* were downregulated by more than 2-fold in Δ10i, suggesting that the reduced expression of multiple genes may contribute to the growth defect of this strain. However, *CMQ270C* expression was comparable to that in WT (Fig. 2B). These results indicate that the growth defect in *Q270*Δi is unlikely to be due to changes in host gene expression (mitochondrial chaperonin hsp60) and supports sieRNA contribution to cell fitness.

**Fig. 2.**
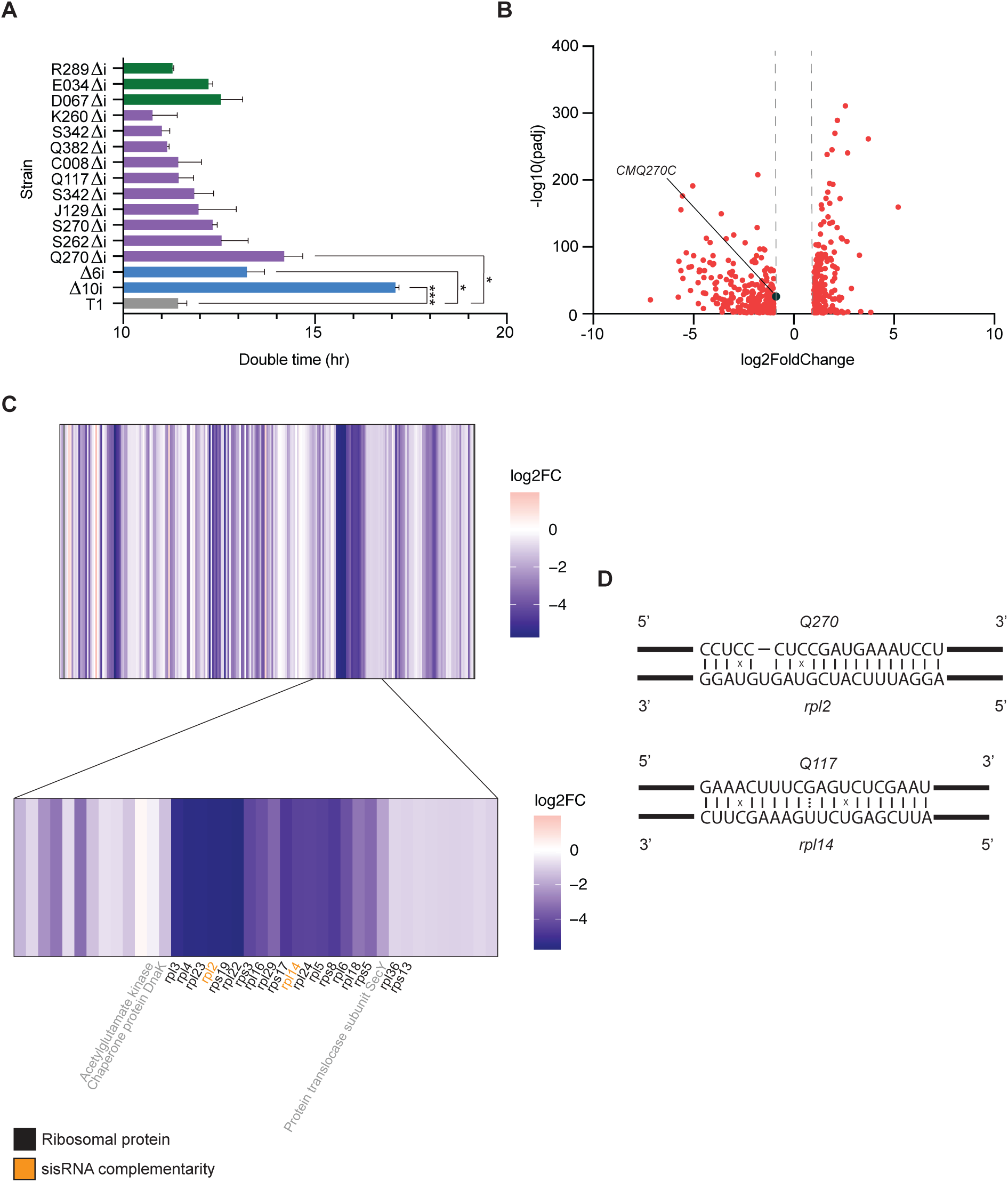
*Q270* may bind and stabilize a chloroplast ribosomal polycistronic locus. **A.** Doubling times of *C. merolae* strains. Cell growth was measured at OD_750_ for 72 hours and taken in biological duplicate. Green, purple, and blue bars represent non-sieRNA-containing introns, sieRNA-containing introns, and multi-intron deletions respectively. Statistical measurements were performed using a paired Student’s T test, where one star represents *P* < 0.05, and three stars represents *P* < 0.001. **B**. Volcano plot of Δ10i differentially expressed genes for log(2) fold change greater than 1 and *P* < 0.05. **C.** Heatmap of chloroplast genes expression in Δ10i along the length of the chromosome. Magnified heatmap on downregulated ribosomal protein cluster. Potential sieRNA targets are highlighted in yellow. **D**. Complementarity of sieRNAs to target ribosomal proteins.

### Ribosomal protein transcripts may be regulated by *Q270* sieRNA

To determine which cellular processes were affected by loss of multiple sieRNAs within the cell, we measured the most differentially expressed genes in the Δ10i strain relative to WT. We found that of the top 30 most downregulated genes, 17 were ribosomal protein genes. When mapped to their chromosomal positions, these genes localized to a single, continuous region of the chloroplast genome (Fig. 2C), indicating coordinated regulation. As plastid genomes can retain prokaryotic features (15), we suspected that this region may be transcribed together as a polycistronic operon.

Consistent with this idea, we identified that intergenic sequences within the cluster were at most 53 nucleotides and in some cases absent. Furthermore, all genes shared the same transcriptional orientation, and we observed full-length transcripts spanning the entire cluster in long-read PacBio sequences. Collectively, these findings support the idea that these ribosomal proteins are transcribed together as a polycistronic operon.

Given that all ribosomal protein genes were downregulated in Δ10i, we hypothesized that sieRNAs are important for maintaining normal expression of the operon. Thus, we assessed the complementarity of sieRNAs to the chloroplast genome. Surprisingly, two sieRNAs, *Q270* and *Q117*, exhibited complementarity to 20-nucleotide regions within the cluster. *Q270* and *Q117* were complementary to the coding sequence of *rpl2,* a gene positioned near the 5’ end of the operon, and *rpl14*, which lies near the middle (Fig. 2D). This observation suggests that *Q270* may be involved in promoting ribosomal protein transcript stability, and loss of the *Q270* sieRNA may reduce ribosome protein expression, contributing to growth defects. Given that *Q117* was identified as a snoRNA, it is possible that it is involved in a different process than *Q270*, which is supported by the lack of growth defect in *Q117*Δi.

### sieRNAs are polyadenylated under heat stress with long poly(A) tails

While *Q270*Δi exhibited slower growth than WT under normal conditions, the other single intron-deletion strains grew at an equivalent or faster rate. We reasoned that if the remaining sieRNAs were functional, they may be important for stress adaptation. In *S. cerevisiae*, excised introns accumulate under nutrient stress and help cells adapt to starvation, while other introns harbour ncRNAs critical to salt stress (1, 16). Given that *C. merolae* is a thermophile, we hypothesized that sieRNAs may accumulate under heat stress conditions.

To test this, we subjected cells to heat stress for one hour and quantified sieRNA abundance using RNA-seq. Surprisingly all sieRNAs appeared upregulated relative to normal conditions (Fig. 3A). We did not detect this response in non-sieRNA containing introns (Fig. 3B). We validated these observations with RT-qPCR, measuring mRNA, pre-mRNA, and sieRNA levels at 57 C, and observed similar changes. (Fig. 3C). However, in contrast to the transcriptomic and RT-qPCR data, Northern blot analysis revealed no change in sieRNA abundance under heat stress (Fig. 3D).

**Fig. 3.**
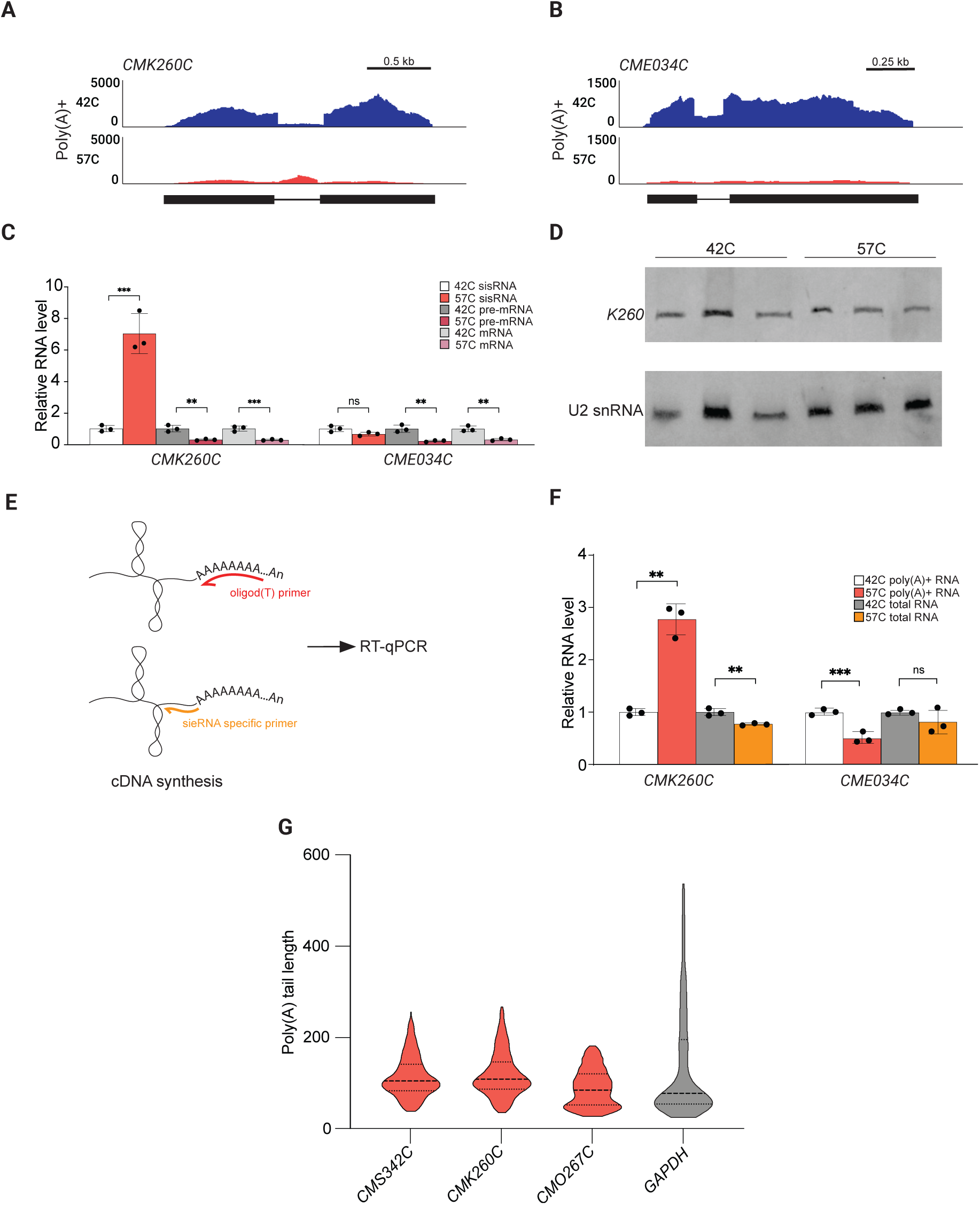
sieRNAs are polyadenylated with long tails under heat stress. **A.** Comparison of *CMK260C* and *CME034C* (**B.**) RNA levels in cells exposed to normal growth conditions (42 C) and under heat stress for one hour (55C). Illumina poly(A) enriched RNA is shown. Experiments were repeated in biological triplicate. **C.** RT-qPCR experiment comparing mRNA, pre-mRNA and sieRNA levels in cells exposed to heat stress at 57 C for one hour to control conditions at 42 C for one hour. *CME034C* was used as a negative intron control. Samples were taken in biological triplicate and compared using a Student’s T-test. **D.** RNA blot comparing the total RNA level of *K260* sieRNA at 42 C and 57 C. *n = 3* for both temperatures. U2 snRNA was probed as a loading control. **E.** Schematic of experiment to test sieRNA polyadenylation using RT-qPCR. **F.** RT-qPCR experiment comparing *K260* sieRNA levels in poly(A) enriched samples to total RNA samples. *CME034C* intron was used as a negative control. **G.** Distribution of sieRNA poly(A) tail lengths at 57 C using an ePAT assay. sieRNA plots are shown in red, while *GAPDH* in grey was used as an mRNA control. *n* = 3 biological replicates.

Given that the RNA-seq analysis was performed on poly(A)-enriched RNAs, and cDNA synthesis targets both polyadenylated and non-polyadenylated RNA, we considered two possibilities. (1) sieRNA accumulation is too small to be measured on an RNA blot despite up to 20-fold increases in the transcriptomic data, or (2) sieRNAs are being polyadenylated in response to heat stress. To distinguish between these possibilities, we used RT-qPCR to compare sieRNA levels under heat stress in poly(A)-enriched cDNA samples to levels detected in total RNA cDNA samples (Fig. 3E). While we measured an increased level of sieRNA in the poly(A)-enriched samples, consistent with the transcriptomic data, total RNA samples had no change or even a small decrease in the sieRNA amplicon under heat stress (Fig. 3F), consistent with the Northern blots. These results are most simply interpreted to mean that sieRNAs are polyadenylated under heat stress.

Notably, some ncRNAs such as rRNAs, snoRNAs and snRNAs acquire short oligo(A) tails during maturation or turnover. These tails are transiently added by the TRAMP complex to facilitate degradation or processing into a ribonucleoprotein complex (17–19). In contrast, mRNA poly(A) tails are longer and promote transcript stability and nuclear export (20,21). Given that sieRNAs were non-coding, we addressed whether their heat-induced polyadenylation matched oligo(A) tail addition in ncRNA control pathways, or if they were more similar to that of mRNAs. To determine this, we measured sieRNA and control mRNA tail length at both 42 C and 57 C using an ePAT assay. At 57 C, sieRNA poly(A) tails were long, ranging from approximately 50 to 200 nucleotides, and matched the length of the control mRNA tails (Fig. 3G). This suggests that sieRNA polyadenylation is not involved in canonical degradation or maturation and may help to stabilize sieRNAs under high heat. Consistent with previous results, we were unable to measure sieRNA polyadenylation under control conditions, supporting the idea that sieRNAs are polyadenylated in response to heat stress.

### sieRNAs are highly structured and may interact with cellular factors

The structure of a ncRNA can influence its interactions and cellular function (22–25). To gain insight into whether sieRNAs interact with other cellular factors, we performed dimethyl sulfate mutational profiling (DMS-MaP) on a subset of sisRNAs under various conditions. This method identifies by their protection from methylation all RNA nucleotides involved in base pairing, or protein binding. As DMS-MaP has not been previously performed in *C. merolae*, we benchmarked the method using U5 snRNA whose structure has been predicted (26). We find that our U5 data closely align with the published structure, validating the accuracy of DMS-MaP in *C. merolae* (Fig 4A).

**Fig. 4.**
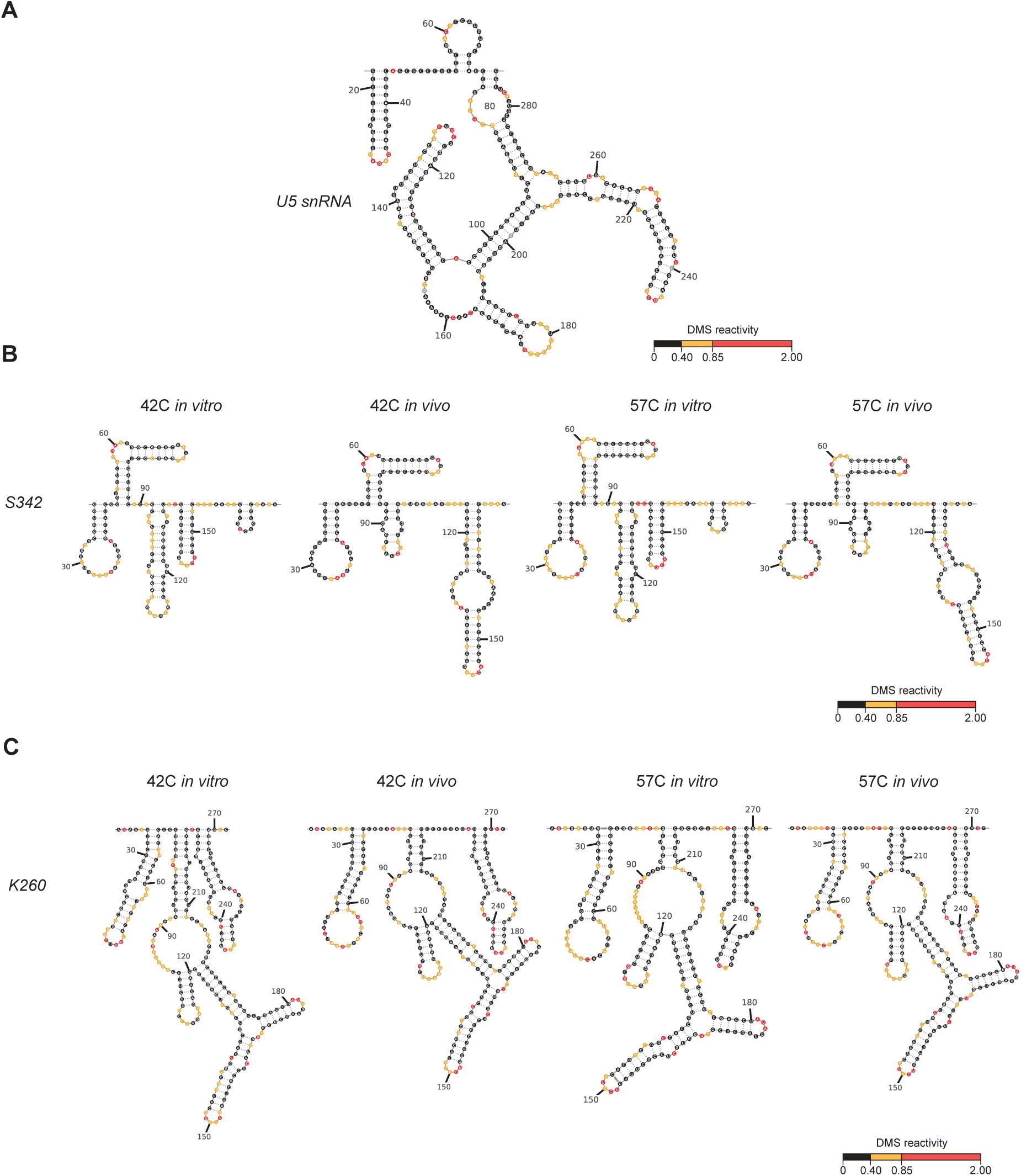
sieRNAs are highly structured and involved in cellular interactions. **A.** Validation of dimethyl sulfate mutational profiling (DMS-MaP) in *C. merolae* using U5 snRNA as a control structure. **B.** Experimental secondary structures of various sieRNAs both *in vivo* and *in vitro* and at normal growth conditions and heat stress. *n* = 1 biological replicates

We reasoned that if sieRNAs were interacting with cellular factors, these contacts would induce differences in DMS reactivity between cell-free and in-cell conditions. For this, we chemically probed sieRNAs *in vivo* and *in vitro.* As sieRNAs are potentially regulated under heat stress, we also compared structures at 42 C and 57 C, to assess possible conformational shifts. Across all conditions, we observed that all sieRNAs were highly structured. We also detected localized shifts in reactivity for all sieRNAs. The second helix of the *S342* sieRNA, for example, was converted into two smaller helices between the 42 C *in vivo* and *in vitro* structures (Fig. 4B). These conformations were consistent at 57 C, indicating that interactions with *S342* sieRNA at 42 C are maintained under heat stress.

We also observed distinct structural features in *K260*. Under normal and heat stress conditions, this sieRNA contained a three-helix junction (Fig. 4C). We also measured increased DMS reactivity *in vitro* within the *K260* three-way junction bulge relative to *in vivo* structures. As this structural element was consistent between conditions, the lower *in vivo* DMS reactivity is unlikely to be due to intramolecular base pairing and could indicate that sequences are occluded due to cellular interactions, for instance protein binding (27,28).

Due to experimental constraints, we were unable to study sieRNAs under 200 nucleotides and over 400 nucleotides long with DMS-MaP. To predict the structure of other sieRNAs, we used a combination of co-variation and thermodynamic structure prediction tools. Similar to DMS-MaP data, computational structures were highly structured and diverse (Fig. S3), supporting previous data that sieRNAs may be involved in regulating various cellular processes.

## DISCUSSION

Despite recent work describing specific functional roles for introns, we lack a clear understanding of why introns are retained through evolutionary time. Given its dramatic intron reduction, we sought to use *C. merolae* as a model to study why the remaining 39 introns were retained, despite evolutionary pressure. We identified that 23 of the 39 remaining introns in *C. merolae* host stable RNAs, which may be important to maintaining cellular function under normal growth conditions and potentially under heat stress. These findings provide an explanation for why these introns have been preserved despite losing nearly 2000 others, and suggest that introns are retained in eukaryotes as a reservoir for regulatory RNAs.

We observed that intronic ncRNAs identified in *C. merolae* are highly conserved in several other thermophilic algal species, while having less retention in mesophilic algae and plants. This suggests that these RNAs could provide an advantage for heat adaptation. Our data suggest that sieRNAs function in various cellular roles.

Interestingly, the *Q270* sieRNA may stabilize a polycistronic mRNA in the chloroplast encoding 17 ribosomal mRNAs by base pairing with *rpl2* transcript. Depletion of this sieRNA results in dramatic downregulation of the cluster and slower growth. This phenotype parallels recent work that found that snoRNAs play an important role in regulating expression of mRNAs (29–31). However, while this work focused on individual transcripts, our work suggests that *Q270* may act as a master regulator for the full polycistronic cluster.

A notable feature of all sieRNAs was their polyadenylation under heat stress. Tail lengths varied from approximately 50 to over 200 nucleotides long. These tails are inconsistent with canonical ncRNA polyadenylation, which drives degradation and maturation, and are more similar to mRNA tail lengths in *C. merolae.* Notably, addition of long, heterogeneous poly(A) tails on rRNA fragments has been observed under heat stress in other organisms. This process, driven by the TENT4b complex, shields against deadenylation to stabilize RNAs and is implicated in driving granule formation (32,33).

As *C. merolae* has been shown to be resistant to heat shock treatments up to 63 C (34), sieRNA poly(A) tails may perform a similar role to drive granule formation and conserve energy. Moreover, heat-induced polyadenylation suggests that sieRNAs collectively serve an alternative function in response to heat stress than their role at 42 C, consistent with a role in adaptation to heat.

Using DMS-MaP and computational analysis, we showed that sieRNAs had conformations suggestive of cellular interactions. We identified conformational shifts between *in vivo* and *in vitro S342* structures, a credible signal of cellular interaction (35,36). However, we also observed that *in vivo* and *in vitro* conformations did not change under heat stress. This may suggest that the *S342* sieRNA is involved in a separate cellular process that has not been explored yet.

Several sieRNAs also contained three helix junctions, a highly stable structural motif that is normally stabilized by tertiary interactions and is a feature of many regulatory RNAs including riboswitches, rRNAs, and RNase P (27,28). Moreover, this motif is characteristic of ligand, protein, or RNA-RNA binding to regulate cellular processes, and may suggest that *K260* sieRNA could share similar regulatory roles (37). Importantly, three-way junctions often form prior to protein interactions, consistent the absence of a structural shift between *K260 in vitro* and *in vivo* conditions (27).

Ultimately, these findings support the theory that introns are important features that host a diverse group of regulatory RNAs important to adapting to their host organism’s ecological niche. As *C. merolae* has lost nearly all of its introns through evolutionary time, sieRNAs help to explain why some introns are retained. The other intronic ncRNAs, snoRNAs and RNase MRP, are predicted to be involved in ribosome biogenesis like the Q270 sieRNA, hinting at the possibility that intronic ncRNAs more generally have been preserved across many lineages for this role. This could explain why introns are so rarely completely lost in extant eukaryotes, despite the overwhelming prevalence of intron loss over evolutionary time (38,39).

## MATERIALS AND METHODS

### C. merolae culturing

The 10D *C. merolae* strain obtained from the Microbial Culture Collection at the National Institute for Environmental Studies in Tsukuba, Japan, was cultured in modified Allen’s autotrophic medium with glycerol (MA2G) in glass graduated cylinders. Six biological replicates were incubated at 42 C with 2% CO_2_ bubbled directly into tubes using an aquarium air pump with 90 μEinsteins of light using a Verilux Happy Light for 24 hours. Cells were collected during log phase growth at an OD_750_ of 1.0-2.0 and then 2 OD units were aliquoted into 1.5 mL microcentrifuge tubes . Three biological replicates were placed in a water bath at 42 C and 57 C for one hour. Cells were then pelleted, and flash frozen with liquid nitrogen to inactivate RNase activity. Pellets were stored at -80 C.

### Total RNA extraction

Cell pellets were resuspended in cold phenol lysis buffer (0.5 M NaCl, 0.2 M Tris HCl (pH 7.5), 0.01 M EDTA, 1% SDS) and sonicated two times for 10 seconds to shear genomic DNA. Three acid phenol/chloroform (pH 4.5) extractions were performed (centrifuged at 15000 x *g* for one minute), followed by two chloroform extractions (centrifuged at 16100 x *g* for one minute) to extract total RNA. For cell extract RNA, one phenol/chloroform (pH 4.5) extraction was performed. Samples were EtOH precipitated and washed in 70% EtOH. RNA was resuspended in ddH_2_O at -80 C until further use.

The concentration and purity of samples were assessed using a Nanodrop One (#ND-ONE-W, Thermofisher Scientific, Waltham). RNA samples with an A_260_/A_280_ ratio between 1.9-2.2 and with an A_260_/A_230_ ratio between 1.8-2.2 were considered clean of cell contaminants and organic solvents. A 1% bleach, 1.5% agarose gel was used with 1 μg of total RNA from each biological replicate to assess RNA quality.

### RNA-sequencing

IsoSeq SMRTbell (Pacific BioSciences) and Illumina RNA-sequencing libraries were prepared as per Scharfen et al. (40). IsoSeq SMRTbell libraries were prepared as recommended by the manufacturer. Full length polyadenylated transcripts were enriched using 500 ng of total RNA while non-polyadenylated transcripts were enriched with 3 μg of total RNA as input. sieRNAs were identified as the longest mapped transcripts within intron boundaries. Illumina libraries were prepared using 1 μg of RNA using TRuSeq Stranded mRNA Sample Prep Kit (Illumina Inc, San Diego, CA) and amplified for 10 cycles. Three replicates were mapped to the *C. merolae* 10D reference genome (8).

### Homologous intron identification

Homologous genes were identified in eight organisms. *C. merolae* sisRNA-containing mRNA sequences were aligned against the genomic sequences of each organism using BLASTX (41). To identify the homologous intron against the sisRNA-containing *C. merolae* introns, TBLASTX from the *C. merolae* mRNA was performed against the genomic sequence of the homologous gene in the eight organisms. Singular gaps at the same position as the intron in the *C. merolae* transcript in the alignment between the mRNA and genomic sequence gaps, were considered to be a homologous intron.

### Northern blotting

Samples were denatured at 65C for 3 minutes in 2X formamide buffer solution and electrophoresed on a 6% polyacrylamide (19:1), 7 M urea gel in 1 X TBE at 400 V. 10 μg of denatured RNA samples were used in biological triplicate and yeast snRNAs were added as a size ladder. RNA was transferred to a Hybond N+ membrane (#RPN303B, GE Healthcare, Chicago, IL) for 35 min at 2.5 mA/cm^2^ using a Panther Semidry Electroblotter (Owl) and then ultraviolet cross-linked. Membranes were pre-hybridized in ULTRAhyb^TM^ – Oligo Buffer (#AM8663, Invitrogen, Waltham, MA) at 42 C for 30 minutes and hybridized at 42 C for 18 hours with probes. Blots were washed (2X SSC, 0.5% SDS) for 30 seconds and then for 30 minutes before being visualized using ChemiDoc MP Imaging System (BioRad). Signals were generated using streptavidin conjugated to IRDye® 800CW (# 926-32230, Li-Cor, Lincoln, NE).

### RT-qPCR validation and analysis

cDNA was synthesized using the iScript Reverse Transcription Supermix (#1708890, BioRad Laboratories, Hercules, CA) and BioRad Reliance Select cDNA synthesis kit (#12012802, BioRad Laboratories, Hercules, CA) according to the manufacturer’s protocol. Oligod(T)s and gene specific primers were used to synthesize cDNA using the Reliance Select cDNA synthesis kit. A no reverse transcriptase (NRT) sample was included for each biological replicate and then pooled together for each temperature.

The annealing temperature of primer pairs were determined using a thermal gradient. Temperatures with the most intense products with the least amount of non-specific binding were chosen. Reaction efficiencies were assessed by performing a standard curve. cDNA stocks were serially diluted 3-fold or 5-fold with at least four serial dilution points in duplicate with iScript cDNA. Standard deviation of duplicate Cq values larger than 0.6 were removed and efficiencies were calculated using CFX Maestro software (BioRad Laboratories, Hercules, CA). Primers with efficiencies between the accepted 90-110% and with R^2^ greater than 0.980 were used for further analysis. Melt peaks generated by CFX Maestro software were used to ensure that a single melt peak was produced from each primer pair, indicating no non-specific binding. A no template control (NTC) was included for each primer pair to ensure minimal primer dimer formation occurred.

Target sisRNAs and reference genes were tested using three biological replicates at 42 C and 57 C of each cDNA type in triplicate on clear 96 well plates (#HSP9601, BioRad Laboratories, Hercules, CA). Fluorescent dye and enzyme were supplied in LUNA 2X Master mix (#M3003E, New England Biolabs, Ipswich, MA). NTC and NRT were run in triplicate for each primer pair. As exon-exon boundaries could not be used to reduce gDNA contamination and sisRNAs contain potential secondary structures, samples with a Cq in NRT or NTC wells lower than 33 were excluded to ensure robust data analysis. To determine if expression was significant (*P*<0.05) between 42 C and 57 C RNA samples, a two-tailed Student’s t-test was performed.

Stability of reference genes were analyzed using GeNorm provided by CFX Maestro software and were kept if M-values were lower than 0.5. Reference genes stability was then analyzed by comparing the change in normalization factor (NF) between biological groups. A two-tailed Student’s t-test was performed to determine if the change in NF was significant.

### ePAT assay

ePAT assay was performed as previously described in Janicke et al. (41). 1 ug of total RNA and 12.5 μM of the universal ePAT anchor or TVN-PAT primer (Table 1) were incubated together at 80 C for 5 minutes and then allowed to cool to room temperature. A mixture containing 5 X SuperScript III buffer (ThermoFisher), 5 mM DTT, 2 mM dNTPs and 5 U of Klenow Polymerase (New England BioLabs) were added to RNA samples and incubated at 37 C for one hour. Enzyme was inactivated at 80 C for 10 minutes and then cooled to 55 C for 1 minute. While still at 55 C, 200 U of SuperScript III (ThermoFischer) was added and then incubated at 55 C for 1 hour. Reverse transcriptase was heat inactivated at 80 C for 10 minutes. cDNA products were then diluted 1:6 and PCR amplified for 33-39 cycles with gene specific forward primers and a tag specific reverse primer (Table 1). Samples were visualized on a 2% agarose gel.

### In-cell DMS treatment and RNA extraction

Falcon tubes were spun at 2000 x *g* for 2 minutes and resuspended in ½ volume folding buffer (300 mM bicine (pH 8.0), MA2G (pH 6.5) media) for a final 2X cell concentration at 42 C or 57 C for 5 minutes. Nine volumes of cells were added to one volume of DMS or neat ethanol, mixed thoroughly, and incubated at initial incubation temperature. For optimization samples at 42 C, 0, 2, 4 and 8M DMS were initially tested then 0, 5 and 6 M were tested. At 57 C, 2-, 3-, 4- and 6-minute incubation periods in DMS were tested.

For experimental samples, 6M for 6 minutes and 6M for 3 minutes were used at 42 C and 57 C respectively. DMS was quenched in 40 volumes of ice-cold 20% (v/v in MA2G) 2-mercaptoethanol and kept on ice for 3 minutes. Samples were spun at 3300 x *g* for 7 minutes. Cell pellets were resuspended in cold phenol lysis buffer (0.5 M NaCl, 0.2 M Tris HCl (pH 7.5), 0.01 M EDTA, 1% SDS) and sonicated for 5 seconds to shear genomic DNA and break down membranes. Two acid phenol/chloroform (pH 4.5) extractions were performed (centrifuged at 17000 x *g* for 15 minutes) followed by one chloroform extraction (16000 x *g* for 2 minutes) to extract total RNA. Samples were EtOH precipitated and washed in 70% EtOH. RNA was resuspended in ddH_2_O and subsequently DNase treated to remove genomic DNA contamination using TURBO DNA-free kit (AM1907, ThermoFisher Scientific, Waltham, MA) according to the manufacturer’s protocol. RNA was then purified using 1.8X volume of MagBind beads (M1378-00, Omega-Biotek, Norcross, GA) according to the manufacturers protocol, except that beads were washed with 70% ethanol 3 times and eluted with water. RNA purity and concentration was quantified using a Take3 nanodrop (Agilent Biotek, Santa Clara, CA).

### Cell-free DMS experiments

RNA was extracted from untreated cells as described above. 2 μg of RNA in 50 μL of water was denatured at 98 C for 1 minute, then snap cooled on ice at 4 C for 1 minute. Equal volumes of cell-free folding buffer (200 mM bicine (pH 8.0), 200 mM potassium acetate (pH 8.0), and 5 mM MgCl_2_) was added and the incubated at 42 C for 20 minutes. Samples were separate into two equal volumes of 45 μL and 5 μL of 1.7 M DMS or neat ethanol was added and incubated at 42 C for 6 minutes. DMS was quenched in an equal volume of 20% 2-mercaptoethanol and RNA was isopropanol precipitated.

### Chemical probing cDNA synthesis and PCR amplification

Reverse transcription was performed as described in Mustoe et al. (23). For in-cell optimization experiments 0.3-1 μg of RNA was used. For experimental samples, 900 ng-1.2 μg of RNA was used as input. Cell-free experiments all used 1 μg of RNA as an input. To optimize DMS reactivity, a one-step PCR approach was performed. One-fifth of purified cDNA products were used as an input [98 °C for 30 s, 30 cycles of (98 °C for 8 s, 66 °C for 20 s, 72 °C for 20 s), and 72 °C for 2 min]. PCR products were purified using 1.8 X volume of beads and quantified using the Take3. Samples were run on a 2% agarose gel at 100 V for 35 minutes and visualized using ChemiDoc MP Imaging System (Biorad Laboratories, Hercules, CA). For experimental samples, a two-step PCR approach was performed. One-fifth of purified cDNA products were used as an input [98 °C for 30 s, 10 cycles of (98 °C for 8 s, 66 °C for 20 s, 72 °C for 20 s), and 72°C for 2 min]. PCR1 products were purified and then one-fifth of product was used as an input [98 °C for 30 s, 20 cycles of (98 °C for 8 s, 65 °C for 20 s, 72 °C for 20 s), and 72°C for 2 min]. PCR2 products were purified using 1.8 X volume of beads. Samples were sequenced using the AVITI sequencing core at UC, Davis. Sequencing reads were aligned using Shapemapper2 software. DMS reactivity was normalized using pairmapper software. Subsequent structure determination was then performed using RNAstructure software and visualized using RNAvigate.

### Homologous structure prediction

A modified version of a pipeline previously performed for viral genome structure detection was employed to detect structure in *C. merolae*’s sisRNAs on the basis of their structural homology with introns in eight other organisms (43). In addition to performing a multiple sequence alignment of the sisRNA with all homologous introns, species homology was confirmed using MAFFT with default parameters (44). The produced MAFFT alignment was also used to inform later steps.

Alignments between sisRNAs and all homologous introns were quite poor with many gaps and isolated stretches of alignment produced. Thus, subsequent analysis was restricted to the strongest alignments, including the homologous introns in *C. yangmingshanensis* and *C. caldarium*.

Alignments were then used as multiple sequence alignment input for four RNA secondary structure prediction programs: RNAz, LocARNA, CaCoFold and KnotAli (45–47). These four utilities were chosen for their relative strengths: RNAz produces additional statistical output including likelihood of conserved structure, LocARNA provides column-wise reliability data, CaCoFold focuses on covariant nucleotide data to predict likelihood of a true conserved structure, while CaCoFold and KnotAli are both capable of predicting pseudoknotted structures. Each of these four utilities were run with default parameters. RNAz was used as a filter; its output SVM RNA-class probability was used as a threshold for whether a sisRNA was capable of forming a conserved RNA structure. Those sisRNAs that achieved a score greater than 0.5 (out of a maximum of 1.0) were deemed capable of forming conserved RNA structure, and analysis with the remaining three utilities was subsequently performed.

