## Supplementary Figure 1 for "Stable, intronic RNAs explain preservation of introns in *Cyanidioschyzon merolae*"

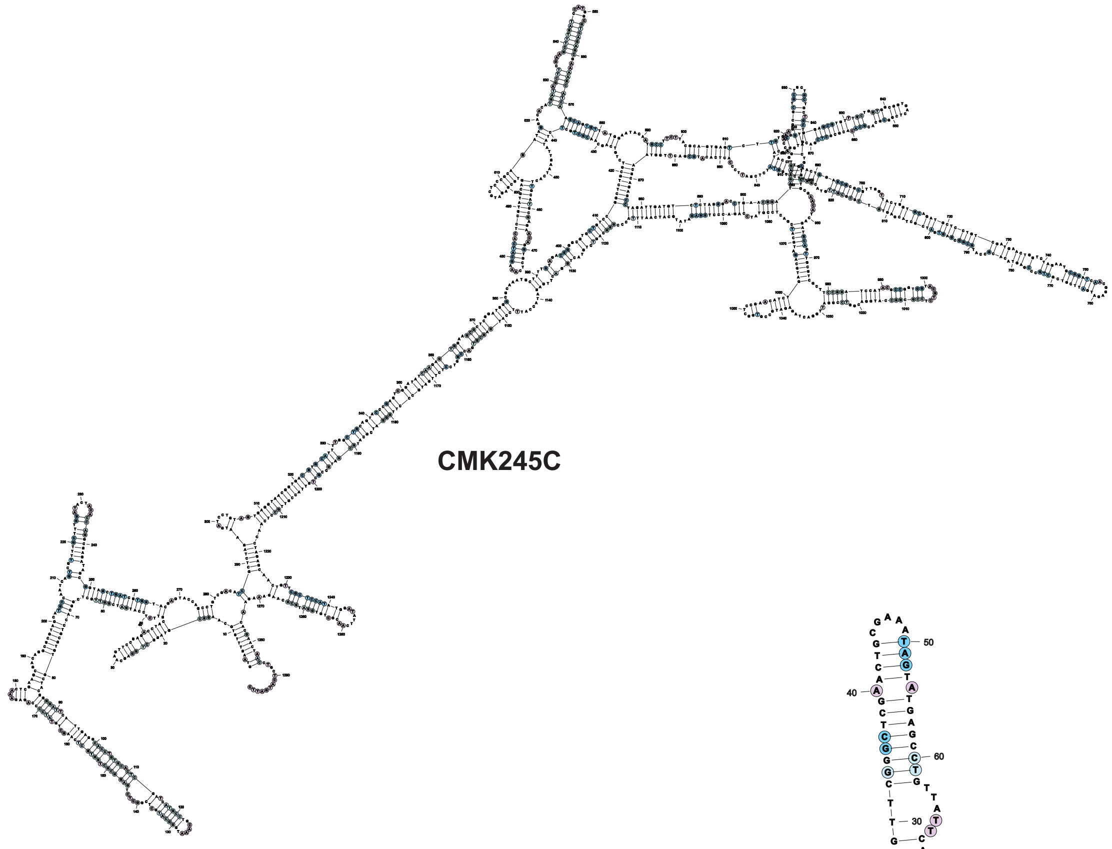

**CMK245C**

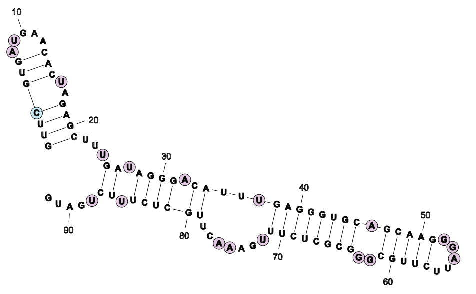

**CMC053C**

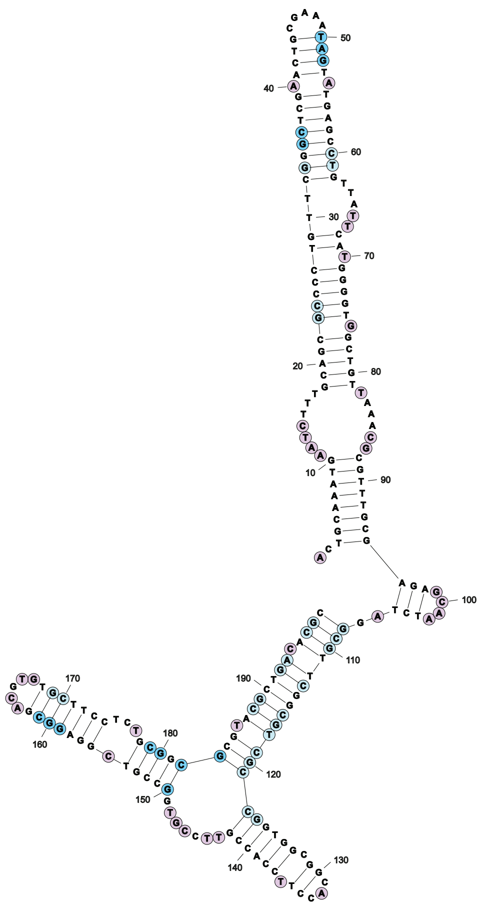

**CMS270C**

**Fig. S1 sieRNAs are structurally diverse.** Predicted structures using RNAz for a subset of sieRNAs.
